# Growth of Stress-Responsive Bacteria in 3D Colonies under Confining Pressure

**DOI:** 10.1101/2024.10.03.616465

**Authors:** Samaneh Rahbar, Farshid Mohammad-Rafiee, Ludger Santen, Reza Shaebani

## Abstract

We numerically study three-dimensional colonies of nonmotile stress-responsive bacteria growing under confining isotropic pressure in a nutrient-rich environment. We develop a novel simulation method to demonstrate how imposing an external pressure leads to a denser aggregate and strengthens the mechanical interactions between bacteria. Unlike rigid confinements that prevent bacterial growth, confining pressure acts as a soft constraint and allows colony expansion with a nearly linear long-term population growth and colony size. Enhancing the mechanosensitivity reduces instantaneous bacterial growth rates and the overall colony size, though its impact is modest compared to pressure for our studied set of biologically relevant parameter values. The doubling time grows exponentially at low mechanosensitivity or pressure in our bacterial growth model. We provide an analytical estimate of the doubling time and develop a population dynamics model consistent with our simulations. Our findings align with previous experimental results for *E. coli* colonies under pressure. Understanding the growth dynamics of stress-responsive bacteria under mechanical stresses provides insight into their adaptive response to varying environmental conditions.

---

Bacteria commonly form colonies on surfaces or in confined natural habitats with porous microstructures upon nutrient availability [1–3]. Although motile bacteria display interesting collective behaviors such as swarming, nonmotile bacterial colonies are abundant in nature and daily life. Within nonmotile colonies, individual bacteria grow, proliferate, and push their neighbors, constructing an evolving network of intercellular mechanical forces across the colony.

Understanding the intriguing spatio-temporal organization of growing bacterial populations has attracted considerable attention over the last few decades because of being of scientific interest and importance in biology, medicine, and technology [4–30]. Structural and morphological complexities arise due to the combined effects of various factors, including mechanical stresses, bacterial characteristics (namely, their shape, growth dynamics, and stiffness), spatial and temporal variations of environmental conditions (such as temperature, viscoelasticity, and nutrient availability), and confinement properties (e.g., size, geometry, stiffness, cell-wall adhesion, etc.).

Mechanical interactions between bacteria develop internal stresses and play a key role in the structural evolution of colonies. Even weak stresses, generated in freely growing colonies in two dimensions, were shown to induce a morphological transition from circular to branched patterns [13]. The interplay of mechanical contact forces and bacterial growth dynamics leads to development of active and passive stresses (align and perpendicular to the cell growth direction, respectively) [14, 15], self-organization of bacteria in ordered subdomains in two dimensions [14– 18], and buckling instabilities which trigger a transition from 2D to 3D structures [19, 20]. Importantly, the imposed mechanical forces on individual bacteria have been found to influence the instantaneous cell growth rate [21, 22]. The mechanosensitivity of bacterial growth affects the length diversity and spatial arrangement of cells, thus, it is expected to influence the stress state of the colony. Stress-responsive bacteria can exploit this stress feedback loop for adaptation to environmental changes. Nevertheless, a detailed understanding of the impact of mechanosensitivity on the dynamics of stress-responsive bacteria and the evolution of structure and stresses across their evolving colonies is still lacking.

Another important factor that influences the structure of bacterial colonies is being constrained to grow against a confinement. The combination of confinement-induced effects and cell stiffness and growth diversifies local cell ordering and self-organized patterns [3, 23–26]. Bacterial natural habitats with rigid microstructures impose a hard constraint preventing the growth of colonies beyond the available spaces. In contrast, a soft constraint— such as being confined with soft agarose pads [20] or growing under a finite confining pressure [27–30]— allows for expansion and further growth of the colony. It has been reported that imposing an external pressure on *E. coli* colonies leads to an unlimited growth of the population, though with a reduced linear rate at long times [27]. How imposing a confining pressure affects the strength of the mechanical interactions between bacteria, the population dynamics, and the evolving size of the colony has remained unexplored so far.

Here we study the growth of 3D nonmotile bacterial colonies under confining isotropic pressure. We consider a model for the mechanosensitivity of bacteria in which the exerted forces parallel to the major axis of stress-responsive bacteria reduce the bacterial instantaneous growth rate. By developing a simulation method to impose a confining isotropic pressure on the bacterial colony, we demonstrate the impact of the pressure on the intercellular forces and address the question of how the population dynamics and the overall size of the colony depend on the confining pressure and mechanosensitivity of bacteria. An analytical estimate is provided for the exponential increase of the cell division time at low mechanosensitivity or pressure, with the pressure dependence being consistent with the experimental results [27]. We also propose a population dynamics model in the presence of confining pressure which accounts for experimental observations [27] and our numerical data.

## Bacterial dynamics under isotropic pressure

We model each bacterium as a spherocylinder with a constant diameter *d*_0_ and a time-dependent length *l*(*t*) of the cylindrical part, i.e. excluding the caps on both ends as shown in Fig. 1(a). The orientation of bacterium *i* is shown by the unit vector 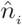 along the major axis of the cell, with components given by 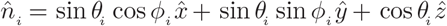 in the lab coordinates (*x, y, z*). The bacteria grow and divide in a three-dimensional space and interact with each other via Hertzian contact forces given by 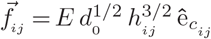 acting from cell *j* on cell *i*, where *E* is the Young’s modulus, *h*_*ij*_ is the overlap distance between the interacting cells, and 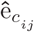 is the unit vector along the line connecting the nearest points on the axes of the *j*th and *i*th cells; see Fig. 1(b).

**FIG. 1.**
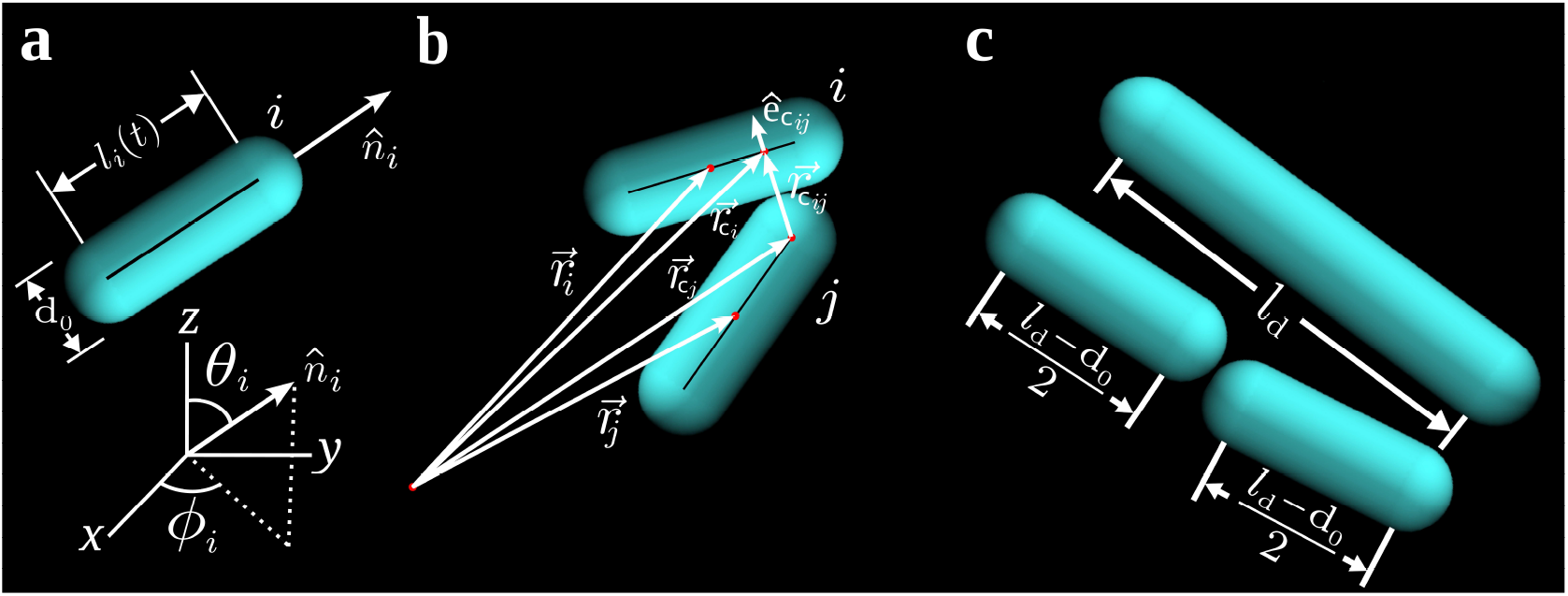
Bacterial model. (a) Geometry of a spherocylindrical cell. (B) A cell-cell contact. (c) A divided cell into two daughter cells after reaching the division length *l*_d_.

To investigate bacterial dynamics in a homogeneous environment, we employ periodic boundary conditions in all directions to eliminate boundary effects. This, however, complicates the application of an external pressure on bacteria in the absence of moving walls or a confining piston. To overcome this problem, we employ a previously developed approach to impose isotropic pressure on a system with periodic boundaries by means of an imaginary piston [31, 32]. In this method, the volume of the system *V* is treated as a dynamical variable whose time evolution is driven by the interplay between a constant external pressure *p*_out_ and the evolving internal pressure *p*_in_ of the system. Denoting the inertia of the piston with

*M*, the change of the volume is governed by

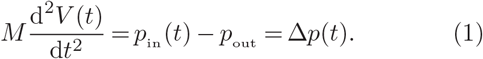

By properly rescaling the momenta and positions of particles due to the volume change, the method was proven to be able to generate isotropic and homogeneous jammed packings of grains [33, 34]. Adapting this method to the overdamped dynamic of bacteria, the change in the position 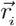 of the *i*th bacterium can be described by

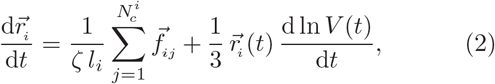

where *ζ* is the drag per unit length and the sum runs over all contacts of the *i*th cell, denoted by 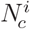. The last term on the right rescales the position according to the relative volume change. Since the imposed isotropic pressure has no influence on the orientation of individual bacteria, the polar and azimuthal angles, *θ*_*i*_ and *ϕ*_*i*_, are independent of *p*_out_ and their time evolution is given by the following equations

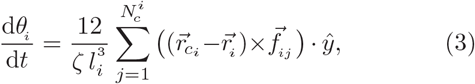

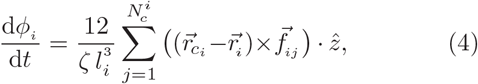

where .,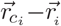 is the contact position vector connecting the center of mass of the *i*th cell to the contact point on the major axis of the *i*th cell.

The gradual growth and division of bacteria and the decrease of the system size due to the imposed external pressure result in the formation of contacts between bacteria. The contacting cells deform each other, which is reproduced in the model by letting them overlap. The elements of the average stress tensor, *σ*_*η,µ*_, can be calculated from the contact forces as [23, 35]

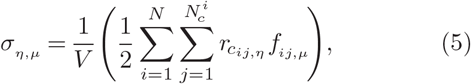

where 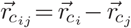 and *N* denotes the number of bacteria. The internal pressure is then given by the trace of the stress tensor divided by the dimension of the system

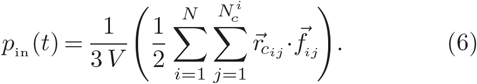

By calculating the pressure difference Δ*p*(*t*) at each time step, we update the volume of the system via Eq. (1), which is required for the position update in Eq. (2). For the integration of equations of motion (2), (3), and (4), we use the implicit first-order Euler scheme. Assuming that an isotropic drag force acts on bacteria, a proper choice for the time step of simulations to generate smooth bacterial rearrangements in the presence of mechanical interactions can be estimated as Δ*t* ≈ *ζ/E*.

In general, the above method enables overdamped dynamics simulations of objects of constant size under confining pressure. To adapt the method to the bacterial dynamics we need to further incorporate the growth and division dynamics of individual bacteria at the beginning of each time step, as described in the following.

### Stress-responsive growth and division model

Cell-cell mechanical interactions cause rearrangements and spatial reorganizations, which influence the overall growth dynamics of bacterial colonies [6, 13, 15, 17, 18, 26, 36]. At the individual cell level, mechanical interactions with the environment have been shown to affect the cell growth dynamics [21, 22, 36]: While the growth rate of the cell can be influenced by the mechanical forces exerted along the major axis of the cell [21], the growth rate is often insensitive to contact forces acting perpendicular to the major axis (except in cases of extremely large forces, which can halt cell growth [36]).

Inspired by these observations, we develop the following stress-responsive model of bacterial growth and division: In the absence of mechanical stresses, a stochastic time-independent growth rate *r*_g,*i*_ is assigned to each bacterium *i*, drawn from a uniform distribution over 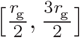 with mean *r*_g_. We model the time evolution of the cell length *l*_*i*_(*t*) under evolving mechanical stresses as

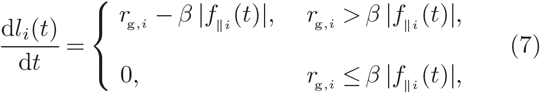

where |*f*_||*i*_ (*t*)| is the total value of the projected forces along the major axis of the *i*th bacterium, i.e. 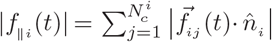. The constant parameter *β* is the mechanosensitivity of the bacterium, characterizing the strength of its response to mechanical stimuli. To take into account the fact that an extremely large lateral force hinders the bacterial growth, we impose an extra constraint on the bacterial growth dynamics: If the overlap distance between the *i*th bacterium and any of its contacting neighbors exceeds the threshold 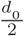 (half of the cell diameter), it pauses the cell growth (i.e. causes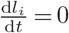) until the overlap decreases below the threshold level again. This constraint has priority over the general growth equation (7).

As the cell length gradually increases according to Eq. (7), it eventually reaches a threshold length *l*_d_ at which the cell division occurs; see Fig. 1(c). Following the division, the two daughter cells tend to inherit the orientation of the mother cell. However, imperfect alignment during division practically leads to a slight stochastic deviation for each daughter cell, which is chosen to be up to maximum 10^°^ in our simulations. Moreover, the growth rates of the daughter cells are also unequal in general, each of them drawn independently from the uniform distribution over 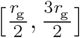 range with the mean value *r*_*g*_.

### Evolution of bacterial colonies

To provide insight into the evolution of colonies of stress-responsive bacteria under confining pressure, we perform simulations based on the bacterial growth, division, and dynamics models described in the previous sections. The default parameter values (unless varied) are taken to be *d*_0_ = 0.5 *µ*m, *r*_g_ = 2 *µ*m.h^−1^, *l*_d_ = 2 *µ*m, *ζ* = 200 Pa.h, *E* = 400 kPa, *M* = 10^−4^ kg.m^−4^, *β* = 0.4 (*µ*m.kPa.h)^−1^, and Δ*t* = 5×10^−4^h. As the simulation starts with the growth of a single elongated bacterium, applying the isotropic confining pressure method leads to a singular behavior of the linear size of the system in different directions. To avoid this technical problem, we do not switch on the confining pressure during the early stages of the simulation, until the bacteria occupy a threshold volume of *V*_c_ = 27 *µ*m^3^ (above which imposing a cubic box shape is definitely feasible). For comparison, the volume of a single bacterium at the onset of division is ∼ 0.46 *µ*m^3^. One can alternatively initiate the simulation with a multicellular random compact colony and equilibrate it before applying the confining pressure.

In a nutrient-rich environment, proliferation of bacteria in the absence of confining pressure results in a freely growing bacterial colony which gradually develops a spherical shape; see Fig. 2(a) and Suppl. Movie. When the confining pressure is imposed according to our developed method, it restricts the growth of the colony within a cubic box with periodic boundary conditions, as shown in Fig. 2(b). We expect that the growth under confinement should lead to denser colonies and affect the population dynamics of stress-responsive bacteria due to the development of internal stresses. In the following, we demonstrate how space-filling and proliferation statistics of the colonies are influenced by the key parameters including the imposed pressure, mechanosensitivity, and growth rate. For ease of comparison, we nondimensionalize the measured quantities using the following units: 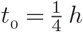, *r*_g0_ = 0.02 *µ*m.h^−1^, *V*_*0*_ = 10^3^ *d*_*0*_ = 125 *µ*m^3^, and 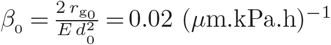.

**FIG. 2.**
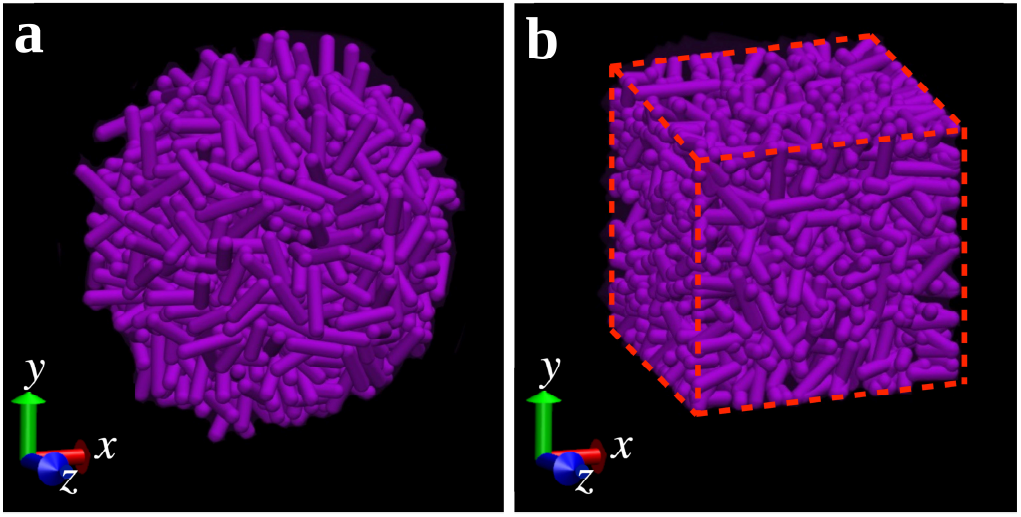
Sample configurations of 3D bacterial colonies growing (a) freely and (b) under confining pressure in a cubic box with periodic boundary conditions.

We first investigate the pressure dependence of the population dynamics. Figure 3 shows the time evolution of the total number of bacteria for two different growth rates and various values of the imposed confining pressure. As explained above, the simulation initially starts with *p*_out_ = 0 until the threshold volume *V*_c_ is reached, which occurs at *t* ∼ 12 *t*_*0*_ or 6 *t*_*0*_ for a bacterial colony with the growth rate of *r*_g0_ or 2 *r*_g0_, respectively. In this initial growth phase, the colonies experience a nearly exponential growth. Note that as a result of the mechanosensitivity of bacteria (i.e. *β* ≠ 0), the instantaneous growth rate of each bacterium varies over time, depending on the exerted local forces which arise due to growth and division dynamics of neighboring bacteria. Switching on the external pressure slows down the population dynamics; an extremely large pressure can even completely prevent the population growth, similar to what happens in a confined geometry with rigid boundaries. A similar pressure dependence of population was reported in experiments on *E. coli* colonies [27]. By imposing the confining pressure, we observe that the slow growth of the total number of bacteria at long times is nearly linear (rather than exponential) with a slope which decreases with increasing *p*_out_. The choice of *r*_g_ influences the results quantitatively but not qualitatively.

**FIG. 3.**
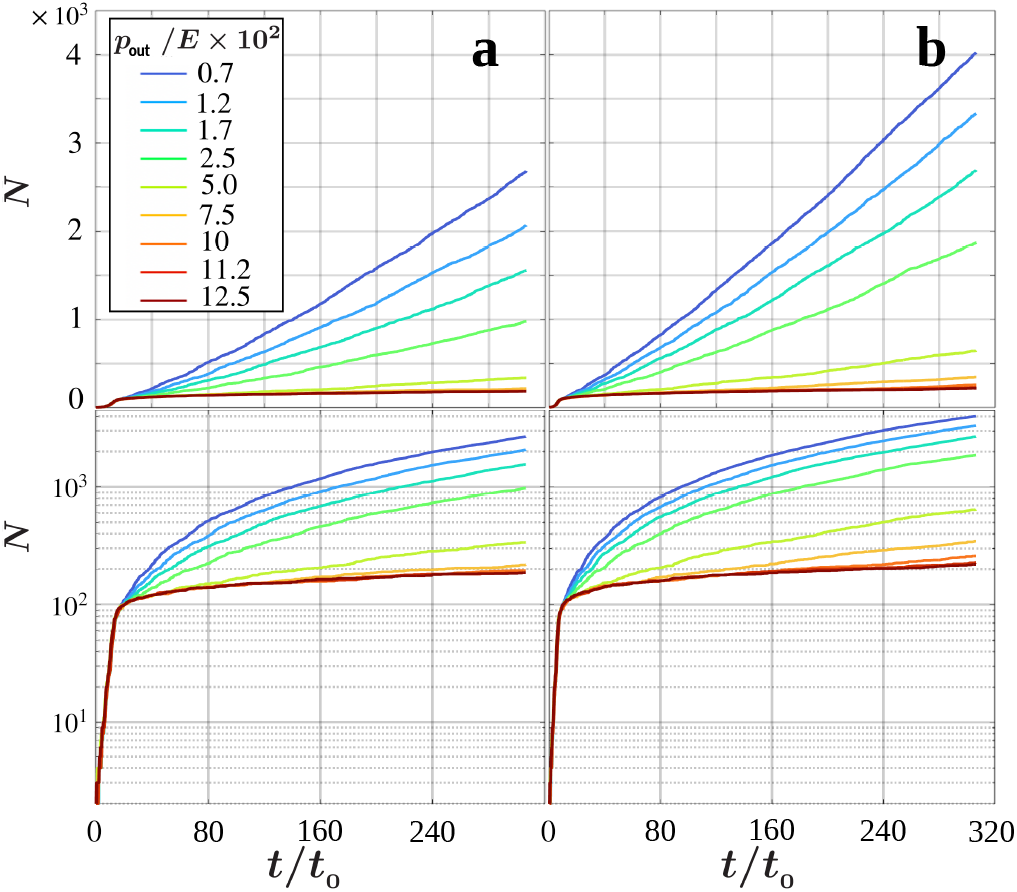
Time evolution of the total number of bacteria *N* for different values of *p*_out_ and (a) *r*_g_ = *r*_g0_ and (b) *r*_g_ = 2 *r*_g0_. The upper and lower panels represent the same plots in linear and log-lin scales, respectively.

The reduction of the population growth rate upon increasing *p*_out_ is expected to influence the evolving volume of the bacterial colony. As shown in Fig. 4, the rate of volume expansion is inversely related to *p*_out_. The growth rate *r*_g_ is also influential; for instance, while an external pressure smaller than 0.07*E* suffices to prevent the expansion of a colony with *r*_g_ = *r*_g0_, a larger pressure *p*_out_ ≥ 0.1*E* is required to stop the expansion of a colony with *r*_g_ = 2 *r*_g0_.

**FIG. 4.**
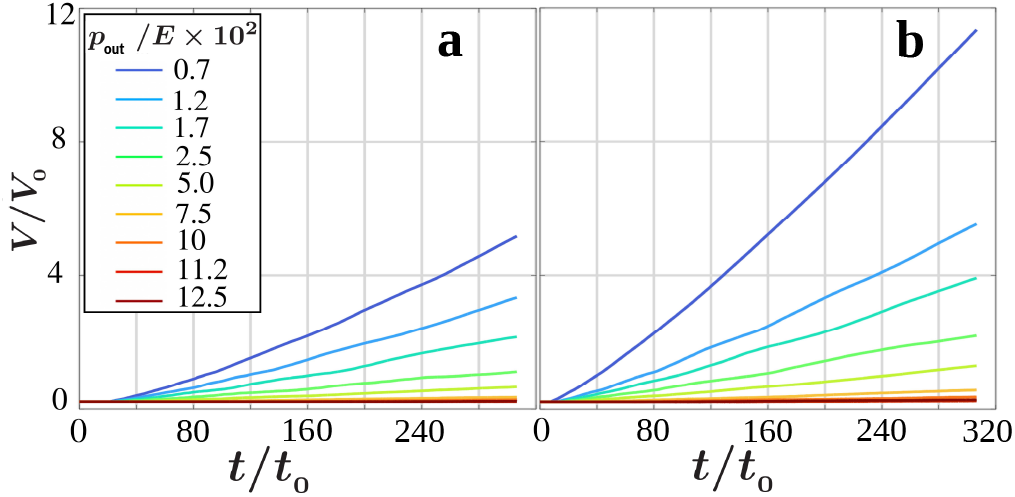
Volume of the bacterial colony as a function of time for different values of *p*_out_ and (a) *r*_g_ = *r*_g0_ and (b) *r*_g_ = 2 *r*_g0_.

The degree of mechanosensitivity of bacteria, reflected in the parameter *β*, determines how far the growth dynamics of individual bacteria and the overall growth of the bacterial colony are affected by the evolving internal stresses. By varying *β* over a wide range, we compare the total number of bacteria *N*_f_ and the volume of the developed colony *V*_f_ after a given long time *t*_f_ ≃ 300 *t*_0_. A larger *β* slows down the growth dynamics of individual bacteria, which slightly reduces the total number of bacteria and the size of the colony; see Fig. 5. For the chosen reference set of biologically relevant parameter values, the sensitivity of the bacterial growth dynamics to the choice of *β* is rather modest: A 10-fold increase in *β* (from *β* = 2*β*_0_ to 20*β*_0_) induces nearly 1% and 3% reduction in *N*_f_ and *V*_f_, respectively.

**FIG. 5.**
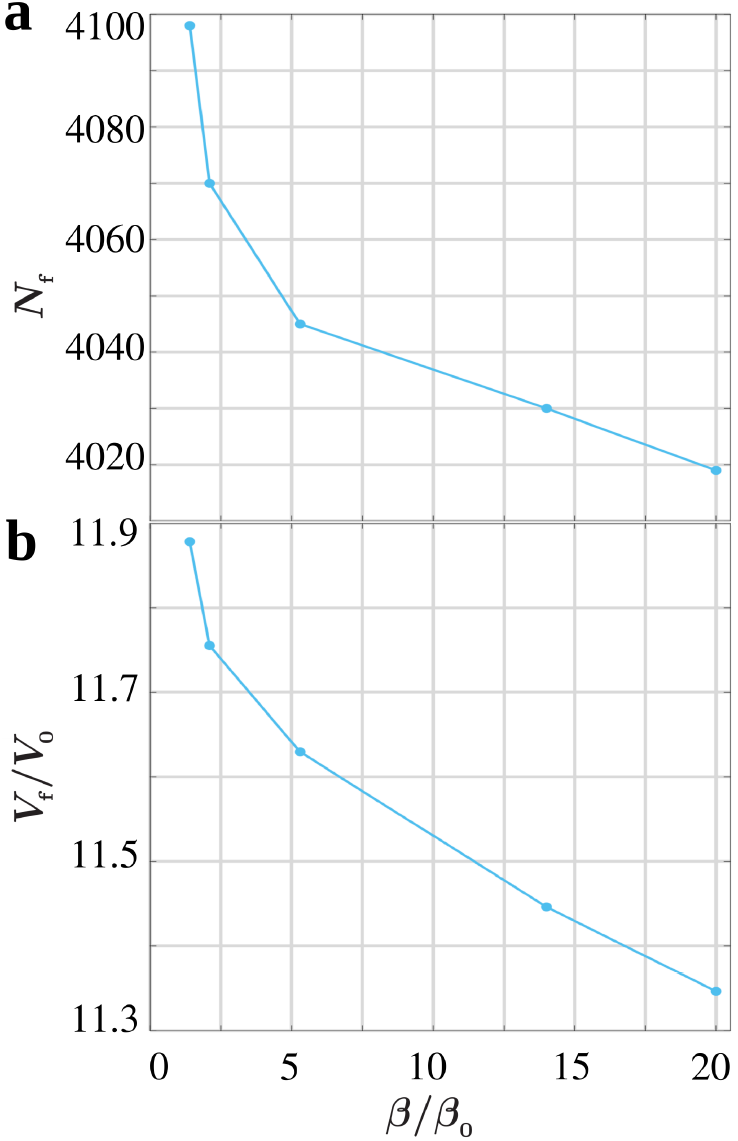
(a) Total number of bacteria and (b) volume of the colony at *t*_f_ ≃ 300 *t*_0_ in terms of the mechanosensitivity *β*.

To better understand the population dynamics under confining pressure, we extract the probability distributions of the overlaps *h* and the instantaneous growth rates 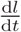 of bacteria across the colony after a long time *t*_f_ ≃ 300 *t*_0_. The increase of the external pressure leads to denser colonies in which the bacteria experience larger deformations. Figure 6(a) confirms that the peak position and mean value of the overlap distribution shift to larger values of *h* upon increasing *p*_out_. Therefore, as the exerted forces on bacteria grow, smaller values of instantaneous growth rates 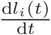 are expected according to Eq. (7). The results shown in Fig. 6(b) reveal that the increase of *p*_out_ leads to smaller instantaneous growth rates of individual bacteria; the probability distribution 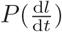 evolves from a broad shape with a mean at intermediate values of 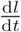 to a monotonically decreasing form with a maximum around 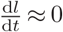. The slower growth rates at larger external pressures lead to longer division times of bacteria. By extracting the division time *t*_d_ via 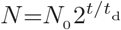, Fig. 7(a) shows that the division time *t*_d_ initially grows with *p*_out_ and eventually plateaus. The increase of *t*_d_ at small *p*_out_ is nearly exponential with a slope which is inversely related to *r*_g_. In previous experiments on *E. coli* colonies [27], an exponential increase in *t*_d_ with *p*_out_ was reported. Interestingly, the slope also decreased with increasing temperature, which is associated with a higher growth rate. As *p*_out_ was increased in these experiments, *t*_d_ eventually diverged because the growth and division dynamics ceased, resulting in an infinite division time for an increasing fraction of bacteria under extremely high stress. In our numerical study, the simulation time window *t*_max_ is finite; thus, we record *t*_d_ ≤ *t*_max_ instead of ∞ for non-growing bacteria, leading to the saturation rather than diverging of *t*_d_ at large *p*_out_. Figure 7(b) reveals that the variation of *t*_d_ versus *β* has a very limited range but follows a similar trend as for *p*_out_ : *t*_d_ grows exponentially with small values of *β* and saturates at large *β*.

**FIG. 6.**
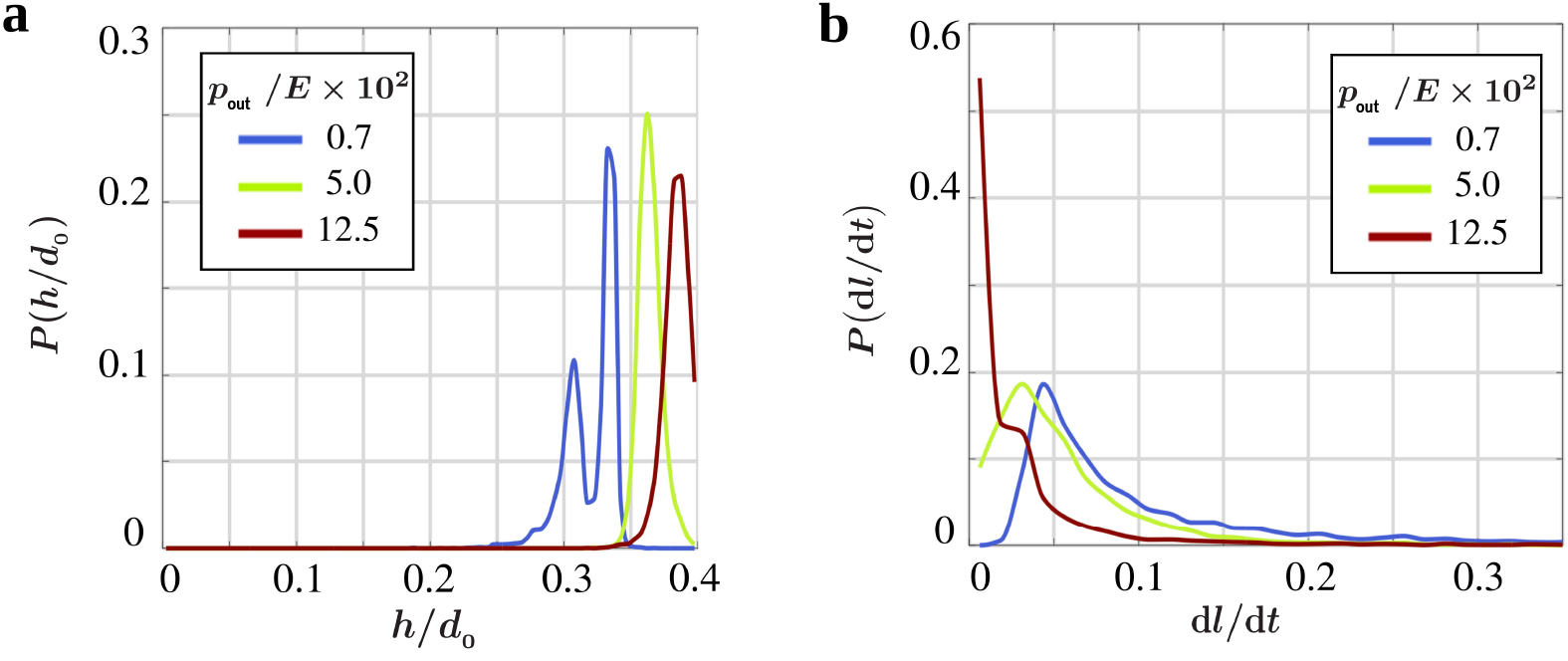
(a) Probability distribution of the scaled overlap *h/d*_0_ and (b) probability distribution of the instantaneous growth rate 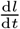 at *t*_*0*_ ≃ 300 *t* for different values of *p*_out_.

**FIG. 7.**
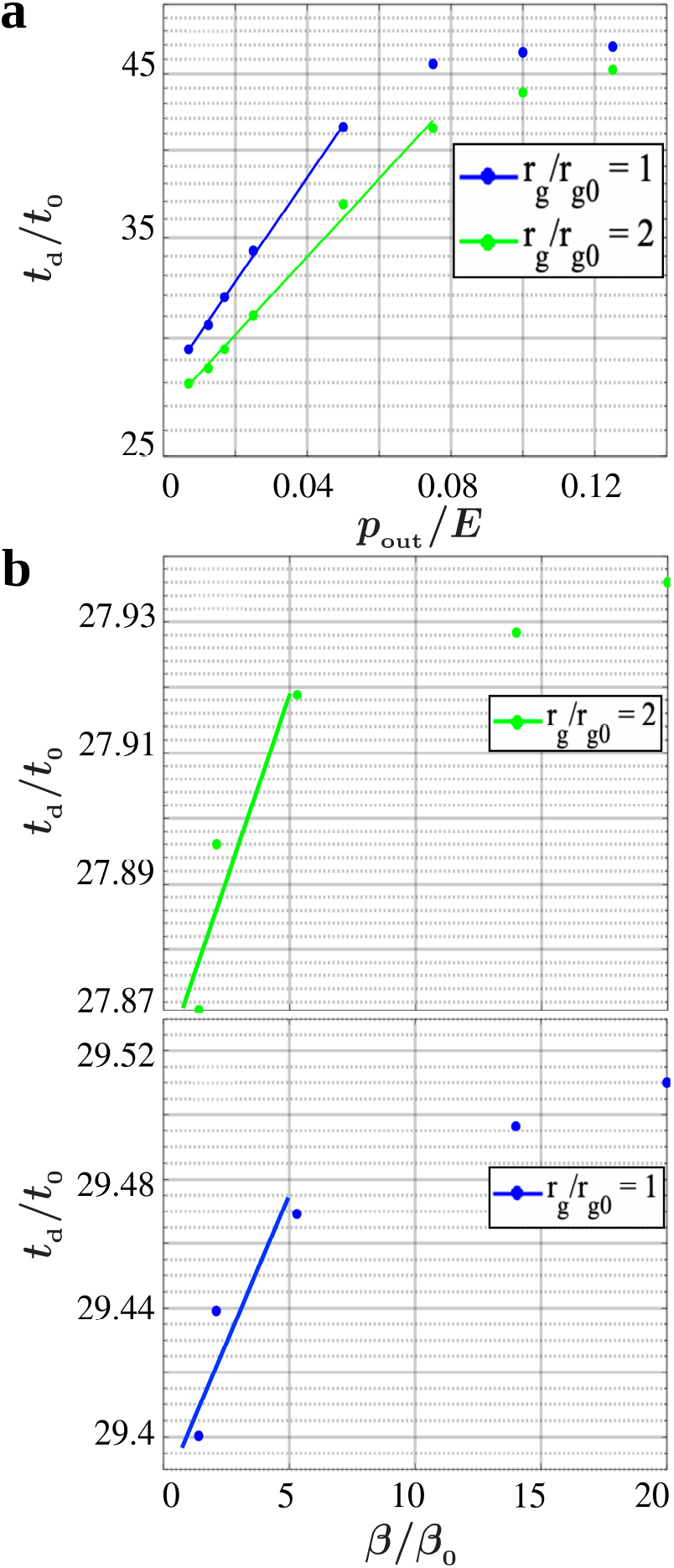
Log-lin plots of doubling time of bacteria *t*_d_ versus (a) *p*_out_ and (b) *β*, for two different growth rates. Other parameters: (a) *β/β*_0_ = 20, (b) *p*_out_ */E* = 0.007. The lines represent exponential fits according to Eq. (11).

### Effective-medium estimate of the doubling time

The doubling time *t*_d_ can be analytically estimated by approximating the fluctuating stress field across the colony at long times (when the internal pressure equals to *p*_out_) by an isotropic homogeneous stress field characterized by *p*_out_. Thus, the discrete contact force network between bacteria is replaced with an effective uniform stress medium. As a result, Eq. (7) for bacterial growth dynamics can be approximated as

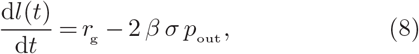

with 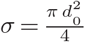 being the cross-section area perpendicular to the major axis of each bacterium. Then, the average length of bacteria at time *t* follows

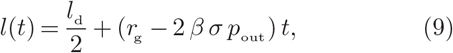

from which the doubling time *t*_d_ can be extracted as

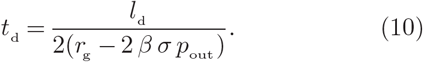

According to this approximation, the doubling time diverges at the threshold confining pressure 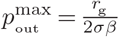 where the mean stresses exerted along the major axis of bacteria reach the required value to fulfill 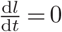. However, even below the 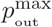 threshold, individual cells in a bacterial colony may randomly experience large forces which prevent their length growth and division (leading to an infinite doubling time). The increase of *p*_out_ enhances the frequency of such stochastic singular events in the system. Therefore, the approximation (10) underestimates *t*_d_ at larger values of *p*_out_ and predicts a pressure threshold 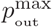 higher than what occurs in practice. As no such singular event occurs in the limit of small *p*_out_, the analytical prediction (10) is expected to successfully capture the behavior. By expanding this equation around *p*_out_ = 0, we can extract an exponential approximation of *t*_d_ at small confining pressures as

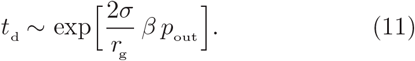

Figure 7(a) shows that the behavior of *t*_d_ at small *p*_out_ is well captured by the exponential relation (11). Expansion of Eq. (10) around *β*= 0 similarly leads to Eq. (11), i.e., *t*_d_ also grows exponentially with *β* in the limit of small *β*. The fit to Eq. (11) shown in Fig. 7(b) verifies that the agreement is satisfactory.

### Population growth model

In order to quantitatively describe the bacterial population growth under confining pressure, we develop a minimal theoretical model for the time evolution of the population of bacteria *N* in the course of internal pressure development. Initially, without the external pressure *p*_out_, the bacterial colony can freely expand and relax the internal stresses by eliminating the deformations induced by bacterial division and growth dynamics. This prevents the increase of the internal pressure *p*_in_ beyond a small minimal level 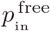 despite the population growth.

Switching on the external pressure 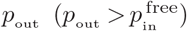 imposes a constraint on the expansion of the colony, resulting in an increase in the number of intercellular interactions in the system and a gradual development of internal pressure. Supposing that the interactions are dominated by binary contacts between neighboring bacteria, our key assumption is that the number of binary interactions and, thus, *p*_in_ grow proportionally with the square of the number of bacteria, i.e. *p*_in_ ∝ *N*^*2*^, for *p*_in_ ≤ *p*_out_. The simulation results shown in Fig. 8(a) support the validity of our assumption: In the initial phase of exponential growth at *p*_out_ = 0, *p*_in_ remains negligible 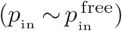 despite the increase of the number of bacteria. By imposing 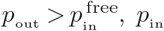 grows nearly linearly with *N* until *p*_in_ and *p*_out_ eventually balance. Beyond this point, the division and growth dynamics of bacteria does not lead to a further increase of *p*_in_ since the system can relax excess internal stresses such that *p*_in_ ≈ *p*_out_ holds while the population can still grow.

**FIG. 8.**
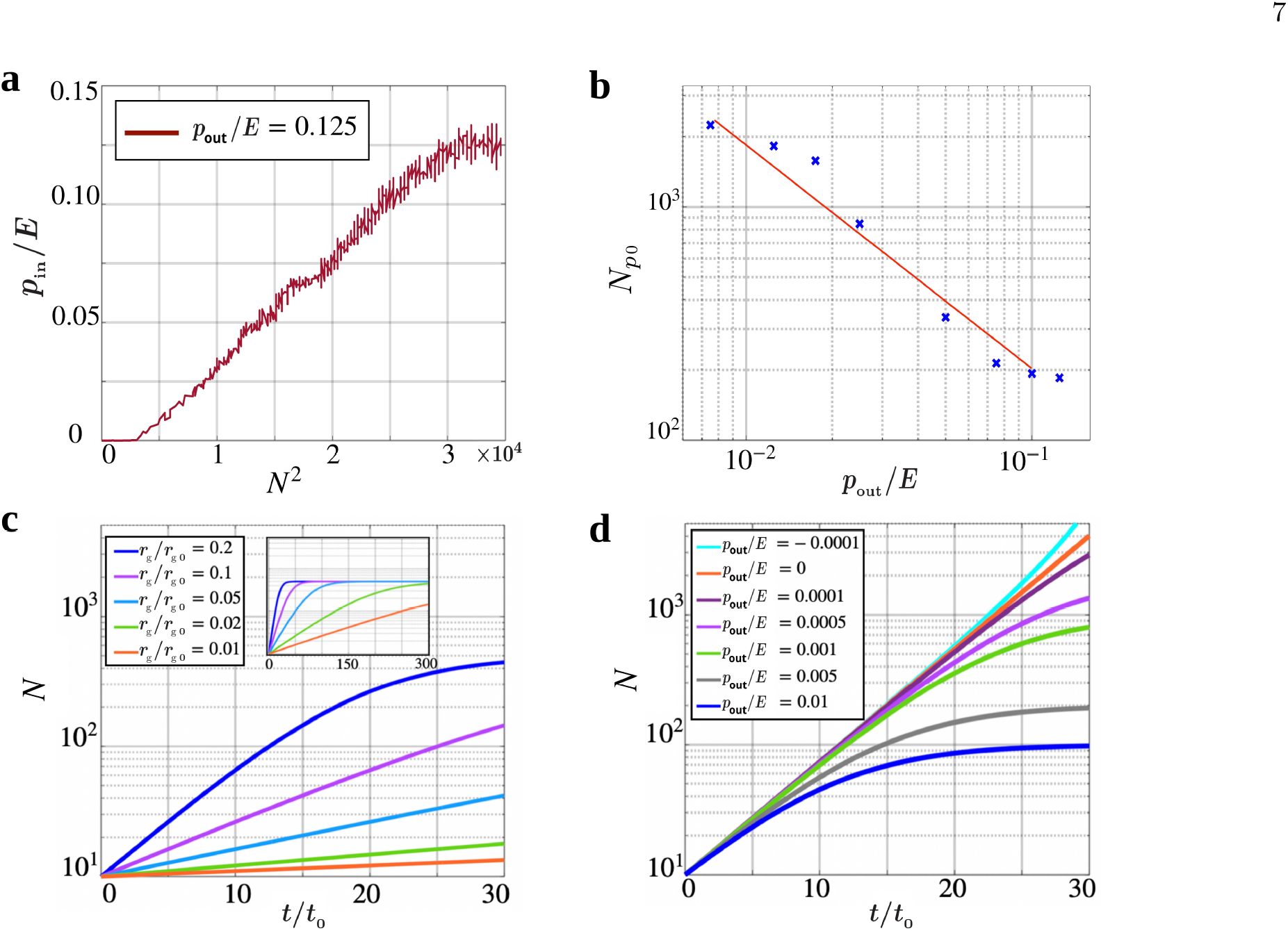
(a) Internal pressure *p*_in_, scaled by the Young’s modulus *E*, versus the square of the increasing number of bacteria at the given external pressure *p*_out_. (b) Population of bacteria *N*_p0_ at the onset of *p*_in_ = *p*_out_ in terms of the external pressure *p*_out_. The line is a fit to *N*_p0_ ∝ 1*/p*_out_. (c),(d) Time evolution of the number of bacteria via Eq. (13) for *N*_0_ = 10 and (c) *p*_out_ */E* = 0.002 and different values of *r*_g_ and (d) *r*_g_ */ r*_g0_ = 0.1 and different values of *p*_out_ */E*. The inset of the panel (c) represents the long time behavior of *N*.

To clarify how the magnitude of *p*_out_ affects the population dynamics, we obtain the population at the onset of *p*_in_ = *p*_out_, denoted with *N*_p_, from the simulations. Figure 8(b) shows that *N*_p0_ decays with *p*_out_ and their relation is roughly captured by *N*_p0_ ∝ 1*/p*_out_ within the studied range of *p*_out_. This leads to our next simplifying assumption that the strength of the imposed constraint is linearly proportional to the applied external pressure.

Based on the above considerations, we propose the following master equation for the time evolution of the number of bacteria

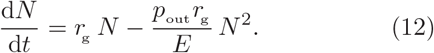

According to the first term on the right-hand side, the rate of change of the population is proportional to the current population leading to an unlimited exponential growth of *N* (*t*). The second term slows the growth rate by increasing the number of binary interactions (∝*N* ^2^) in the presence of external pressure. Equation (12) implies that the evolution of *N* is entirely determined by the confining external pressure, Young’s modulus, and mean growth rate. The solution of this logistic growth model is given by

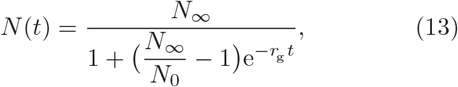

with *N*_∞_ = *E/p*_out_ being the carrying capacity of the system and *N*_0_ the initial number of bacteria when *p*_out_ is switched on.

Figures 8(c),(d) represent the time evolution of *N* via Eq. (13) for different values of mean growth rate and external pressure. It can be seen that our simple model qualitatively reproduces the observed behavior in simulations (see the time evolution of *N* after the initial exponential growth phase in lower panels of Fig. 3 for comparison). Note that choosing *p*_out_ = 0 reduces Eq. (12) to a simple exponential growth and a negative external pressure *p*_out_ *<* 0 even accelerates the population growth, as shown in Fig. 8(d).

The asymptotic behavior of *N* (*t*) is shown in the inset of Fig. 8(c), highlighting that *N* (*t*) reaches a plateau at long times. This behavior differs from the simulation results in Fig. 3, where *N* (*t*) continues to grow at long times with a slope which is inversely related to the external pressure. In the logistic growth model Eq. (12), the maximum possible number of bacteria *N*_∞_ is set by the ratio *E/p*_out_, thus, the carrying capacity is fixed. Nevertheless, as we previously showed in Fig. 8(a), the carrying capacity of the system grows forever beyond the onset of *p*_in_ = *p*_out_ since the system can expand and relax the excess internal pressure generated by the bacterial division and growth dynamics. To consider this effect in our model, we rewrite Eq. (12) such that a linearly growing term with time, 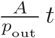, is added to the maximum possible number of bacteria (with *A* being a constant) to allow the gradual increase of the capacity of the system. Therefore, we propose the following master equation for the time evolution of the number of bacteria in the colony under confining pressure

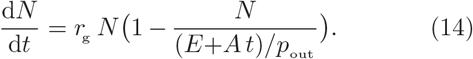

This equation can be numerically solved for a given set of {*r*_g_, *p*_out_, *E, A*} values to obtain *N* (*t*). The results shown in Fig. 9 reveal that the asymptotic continuous growth of *N* (*t*) is reproduced with the modified master equation (14) and even the inverse *p*_out_ -dependence of the slope at long times in captured.

**FIG. 9.**
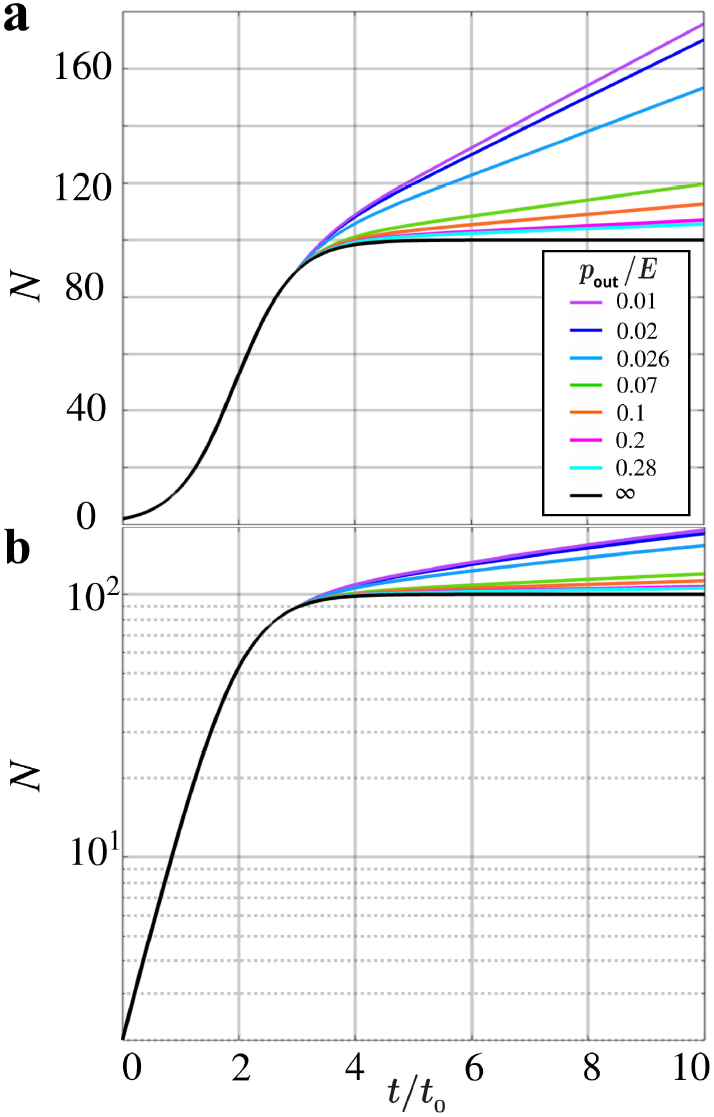
Time evolution of *N* according to Eq. (14) in (a) linear and (b) log-lin scales for *N*_0_ = 2, *r*_g_ */ r*_g0_ = 1, *A/E* = 10^−4^ h^−1^, and different values of the external pressure.

### Discussion and conclusion

We have numerically studied the evolution of colonies of stress-responsive bacterial under confining isotropic pressure. To generate homogeneous colonies, boundary effects have been eliminated by employing periodic boundary conditions, which makes the imposing of a confining pressure challenging. We have implemented a method based on rescaling the momenta and positions of cells, and adapted it to the overdamped dynamics of bacteria. The growth dynamics of the colony is influenced by the imposed pressure through affecting the intercellular interactions as well as the growth dynamics of individual stress-responsive bacteria. By introducing the mechanosensitivity in our model, the sensitivity of the bacterial growth to the exerted stresses can be tuned. The validity of our model is ensured by the remarkable agreement between our numerical predictions and experimental results of evolving *E. coli* colonies under confining pressure [27]. The simulation method presented in this work can be efficiently parallelized for large-scale simulations of bacterial systems since the varying volume method is compatible with effective parallelization techniques based on adaptive hierarchical domain decomposition with dynamic load balancing [37]. The simulation method can be also straightforwardly generalized to model assemblies of motile cells.

In the present work we have chosen constant values for the model parameters for simplicity. This includes the structural and mechanical properties of bacteria (such as cell diameter *d*_0_ and Young’s modulus *E*), bacterial growth parameters (i.e. growth rate *r*_g_ and division length *l*_d_), and environmental properties including the drag per unit length *ζ* and temperature (implicitly). Nevertheless, this does not reflect the diverse phenotypic characteristics of bacteria such as their morphology and differentiation variability. Real distributions of the relevant quantities can be extracted experimentally and served as input for simulations to produce quantitatively comparable statistics. To this end, the mechanical interactions and equations of motion in our model would require slight modifications.

Introducing mechanosensitivity in our model can have complex effects on the structure and dynamics of bacterial colonies. For the chosen set of parameter values in our study, the sensitivity of the results to the choice of *β* has been modest. However, more generally, the reduction of the growth rates of mechanosensitive bacteria under mechanical stresses influences their length diversity and spatial rearrangements, which changes the stress state of the system. Such a feedback loop can be exploited by stress-responsive bacteria to adapt themselves to environmental changes. It will be interesting to explore the impact of mechanosensitivity on structure, dynamics, and adaptation of bacterial colonies for broader biologically-relevant ranges of model parameters.

In the absence of quantitative experimental data, we have chosen a simple linear force-dependence form for the mechanosensitivity in Eq. (7). This functionality can be adapted to the specific response of each different type of bacteria to the imposed stresses. Stress-responsive bacteria in our model experience time-dependent instantaneous growth rates due to the variations of mechanical stresses. The model can be extended to take into account the spatial and temporal changes of growth rates in response to local nutrient availability [10, 38] or viscoelasticity variations [39]. According to experimental observations [21, 27, 36], imposed stresses can additionally affect the threshold division length and even shape of bacteria and induce aging of cell mechanical properties. Algorithmic implementation of these concepts requires further quantitative experimental data.

To summarize, we have developed methods and carried numerical simulations to study evolving 3D colonies of stress-responsive bacteria under confining pressure. Our results demonstrate how physical interactions regulate biological processes at the microscale and highlight the intricate feedback mechanism between mechanical stimuli and bacterial growth dynamics. Understanding the interplay of mechanosensitivity, structural characteristics of bacteria, and mechanical interactions can provide insights into the adaptive responses under varying environmental conditions and may inspire novel approaches to control bacterial infections.

This work was supported by the Deutsche Forschungsgemeinschaft (DFG) within the collaborative research center SFB 1027 and also via grants INST 256/539-1, which funded the computing resources at Saarland University. R.S. acknowledges support by the Young Investigator Grant of Saarland University, Grant No. 7410110401.

## Notes

### Competing Interest Statement

The authors have declared no competing interest.

